# MesoTIRF: a Total Internal Reflection Fluorescence illuminator for axial super-resolution membrane imaging at the mesoscale

**DOI:** 10.1101/2022.08.19.504513

**Authors:** S. Foylan, W. B. Amos, J. Dempster, L. Kölln, C. G. Hansen, M. Shaw, G. McConnell

**Author notes:** Electronic mail.

## Abstract

Total Internal Reflection Fluorescence (TIRF) illumination bypasses the axial diffraction limit of light by using an evanescent field to excite fluorophores close to a sample substrate. TIRF illumination significantly improves image contrast, allowing researchers to study membrane structure and dynamics with localized reductions in photobleaching. However, a significant limitation of most TIRF microscopes is the relatively small field of view (FOV). TIRF objectives require a high numerical aperture (NA) to generate the evanescent wave. Such lenses invariably have a high magnification and result in a ∼ 50 µm diameter imaging field, requiring many subsequent images for accurate statistical analysis. Waveguide and prism-based TIRF systems are, in principle, compatible with lower magnification lenses to widen the FOV but these have a correspondingly low NA and lateral resolution. To overcome these limitations, we present a prism-based TIRF illuminator for the Mesolens - a specialist objective lens with the unusual combination of low magnification and high NA. This new imaging mode - MesoTIRF - enables TIRF imaging across a 4.4 mm x 3.0 mm FOV. We demonstrate evanescent wave illumination of cell specimens, and show the multi-wavelength capability of the modality across more than 700 cells in a single image. MesoTIRF images have up to a 6-fold improvement in signal-to-background ratio compared to widefield epi-fluorescence illumination, and we illustrate the benefit of this improved contrast for the detection and quantification of focal adhesions in fixed cells. Fluorescence intensities and resolvable structural detail do not vary considerably in homogeneity across the MesoTIRF FOV.

## I. INTRODUCTION

Total Internal Reflection Fluorescence (TIRF) microscopy is an established imaging technique in cell biology^1^. It relies on delivering excitation light such that it is incident on a refractive index boundary at the microscope specimen plane at a super-critical angle. The subsequent Total Internal Reflection (TIR) results in a rapidly decaying evanescent field which penetrates to a depth on the order of several hundred nanometres. Such illumination allows structure below the axial diffraction limit to be visualized with high contrast, while minimizing photobleaching of the specimen by reducing the illumination volume. TIRF microscopy has been used extensively in cell biology to image cell contacts^1^, study cell adhesion^2,3,4^, ion channels^5^, endocytosis^6^ and the self-assembly of filamentous proteins^7^. Generating TIR at the specimen plane can be achieved using a TIRF objective, a waveguide^8^, or a prism^9^. To support super-critical illumination, TIRF objectives have a numerical aperture (NA) between 1.45-1.5. However, due to their limited size, these objective lenses have an associated high magnification, typically 60x or 100x. As such, the field of view (FOV) is restricted to around 50 µm x 50 µm and the microscope can only capture a few cells in a single image^10^. Stitching and tiling methods can be used to image larger specimen areas, but these methods are time-consuming and routinely introduce artefacts into the resultant image. Waveguide-based TIRF obviates the need for a high NA objective lens allowing images of up to 0.5 mm in diameter containing tens of cells using custom-designed chips^10,11^. Prism-based TIRF uses off-the-shelf components and is therefore lower cost and easier to implement than other methods, and, like waveguide TIRF, is compatible with any objective lens. This potentially allows for larger FOV imaging than possible with TIRF objectives. However, for both waveguide and prism-based TIRF, the detection objective lens remains a fundamental limitation when considering FOV. Low magnification objective lenses that support wide FOV imaging typically have a low NA, which in turn leads to low resolution images.

Here, we report MesoTIRF - prism-based TIRF microscopy using the Mesolens for imaging over a large FOV with high lateral and axial resolution. The Mesolens^12^ combines low magnification with a high numerical aperture (4x/0.47 NA), and our dual-wavelength MesoTIRF illuminator generates an evanescent wave to exploit its full FOV (4.4 mm x 3.0 mm). We present details of the optical setup and show the utility of MesoTIRF for high-contrast, high-resolution imaging of fluorescently-labeled proteins in fixed cells.

## II. METHODS

A schematic diagram of the optical set-up is shown in Figure 1. The illumination laser source was a tunable wavelength (Chameleon Ultra II, Coherent) Titanium Sapphire laser pumping an optical parametric oscillator (OPO) (Compact OPO-Vis, Coherent). The second harmonic of the signal wavelength output of the OPO was used as the laser source, with 500 nm and 585 nm selected for dual-wavelength TIRF imaging. This choice of wavelengths was informed by the specification of the custom 100 mm diameter Pinkel-type^13^ chromatic reflector and barrier filters used for Mesolens imaging which allows fluorophore excitation/emission combinations of 505 nm ± 25 nm/542.5 ± 7.5 nm and 575.5 ± 22.5 nm/677.5 ± 72.5 nm. For comparison of the performance of the MesoTIRF illuminator with widefield epi-fluorescence illumination (WF epi) we used 504 nm (bandwidth = 19.4 nm) and 584 nm (bandwidth = 27 nm) light emitting diodes for wide-field illumination (pE-4000, CoolLED).

**Figure 1:**
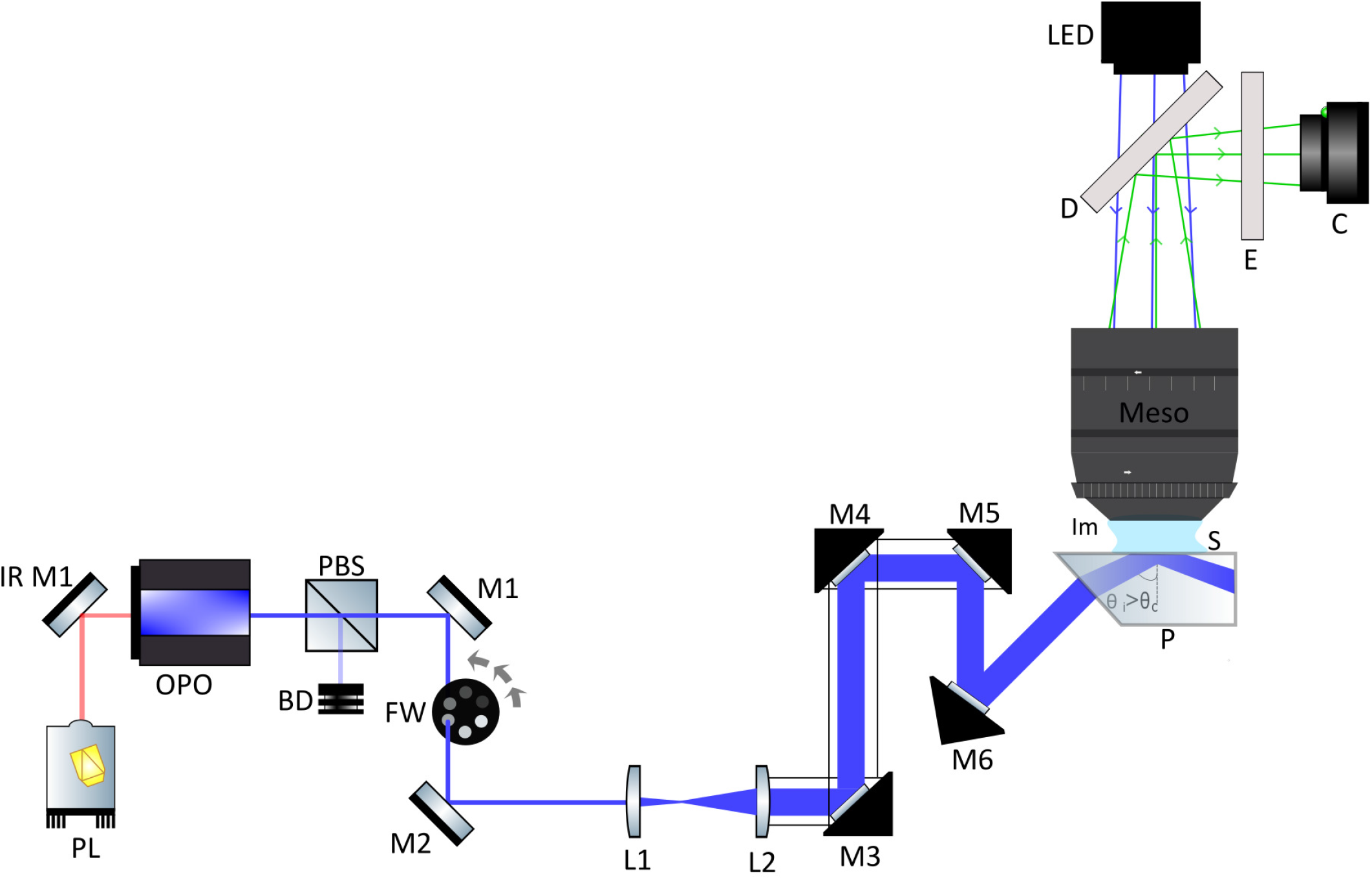
Schematic of MesoTIRF: PL: Titanium Sapphire pump laser (Ultra II, Coherent), IR M1: Infrared mirror (BB1-E03, Thorlabs), OPO: Optical Parametric Oscillator (Chameleon OPO-Vis, Coherent), PBS: polarising beam splitter (CCMS-PBS201/M, Thorlabs), BD: beam dump, M1-6: visible broadband dielectric mirrors (Thorlabs, BB1-E01), FW: filter wheel with 5 neutral density filters (Thorlabs, FW1A), L1: 50 mm planoconvex lens (LA1131-A-ML, Thorlabs), L2: 100 mm planoconvex lens (LA1509-A-ML, Thorlabs), M3-6 mounted in right angled cage mounts (KCB1C/M, Thorlabs), P:45° borosilicate glass prism (Mesolens Ltd.), S: sample, Im: immersion fluid (distilled water), Meso: Mesolens objective element (^12^), D: dichroic filter & E: emission filter(custom from Chroma), C: chip-shifting camera sensor (VNP-29MC; Vieworks), LED: 504 nm and 584 nm LEDs from LED module (pE-4000, CoolLED)

The optical power of the OPO output beams at the specimen plane were adjusted using a combination of a polarizing beamsplitter cube and a variable neutral density filter wheel. Next, the beam was expanded by a Keplerian telescope consisting of 50 mm and 100 mm focal length plano-convex lenses (anti-reflection coated for 350 - 700 nm). A first surface reflector in a kinematic mount was used to adjust the angle of incidence of the beam to 86° at the top surface of a 45° borosilicate glass dove prism which served as the MesoTIRF prism. The theoretical evanescent field depth for a wavelength of 504 nm in a borosilicate prism (n=1.51) at this incidence angle is 56 nm ^14^. The 25 mm thick prism has a top surface of 20 mm by 70 mm, and was placed on top of a computer-controlled specimen stage (ProScan III, Prior Scientific) for accurate positioning of the prism in three dimensions. To capture the large, high-resolution images produced by MesoTIRF we used a chip-shifting camera sensor (VNP-29MC; Vieworks) which recorded images by shifting a 29-megapixel CCD chip in a 3 × 3 array^15^. Reconstruction of each image (260 Megapixels, 506 MB) took approximately 5 s on a typical computer workstation.

To confirm evanescent illumination, we prepared a specimen of murine fibroblast cells (3T3-L1) labeled with both a fluorescent nuclear marker (SYTO Green) and an antibody labeling against paxillin, a focal adhesion component^16^. We hypothesized that the fluorescent emission from the nuclear marker would be visible in WF epi but with MesoTIRF the cell nucleus would be too far above the basal membrane to be excited by the evanescent wave. To image the SYTO Green stain, the camera exposure time and the camera gain were set to 2 s and 1 X, respectively for both WF epi and MesoTIRF. In WF epi, the 504 nm LED power was adjusted to 25 mW to excite fluorescence from the stained nuclei without saturation. For MesoTIRF, the maximum available laser power at the specimen plane of 3.48 mW was used at a wavelength of 500 nm. To image the antibody against paxillin that was conjugated to Alexa Fluor Plus 594, the camera exposure time and camera gain were set to 2 s and 70 X, respectively for both WF epi and MesoTIRF. The optical powers of the 584 nm LED (22 mW) and the 585 nm laser (3.48 mW) were adjusted to produce images of the fluorescently labeled paxillin with a similar fluorescence signal intensity. To estimate the number of cells, the ‘Surfaces’ model in Imaris (Imaris 9.8, Oxford Instruments) was used for object detection of SYTO Green labeled nuclei in the WF epi image.

A comparison of signal-to-background ratio (SBR) in WF epi and MesoTIRF from this dual labeled sample was obtained by taking line intensity profiles in ImageJ^17^ through focal adhesions in the images from each modality. Following transfer of this data to Python, the peaks of each focal adhesion where detected using the find_peaks() function in the SciPy^18^ library and scaled against the minimum signal intensity.

To evaluate the capability of MesoTIRF for dual-wavelength imaging including an assessment of the uniformity of illumination, a fixed HeLa cell specimen was prepared, using an anti-paxillin antibody that was conjugated to Alexa Fluor Plus 594 and fluorescein phalloidin which stains F-actin. For each sequential image, an exposure of 2 s and a camera gain of 30 X was used.

To evaluate the improvement in SBR in MesoTIRF compared with WF epi, the same specimen was imaged but only the anti-paxillin conjugated to Alexa Fluor Plus 594 was excited. To measure the SBR Trainable Weka^19^ was used to identify and segment objects from both WF epi and MesoTIRF images. A selection mask was extracted from this segmentation, which enabled us to derive the mean detected signal in a cell from the raw images using ImageJ^17^. The background signal was calculated for each image by selecting three background regions of interest, calculating the mean intensity, and averaging these measurements. This workflow was carried out for six ROIs separated by at least 0.5 mm across the full FOV to evaluate variation in SBR, and the uniformity of illumination.

To demonstrate MesoTIRF in another cell type, the human mesothelial cell line MeT-5A was fixed and labeled with antibodies against paxillin and tubulin, which were conjugated to Alexa Fluor 488 and Alexa Fluor Plus 594 respectively. These (adhesion and cytoskeletal) proteins are within the reach of an evanescent field. This data set is included as Supplementary Information.

Cell culture, fluorescent labeling and specimen preparation methods are included as Supplementary Information.

## III. RESULTS

Figure 2 shows a comparison of WF epi with MesoTIRF images of dual-labeled fixed 3T3-L1 cells prepared with fluorescent staining for nuclei (green) and paxillin (magenta). The full FOV WF epi image is shown in 2A, with a region of interest (ROI) indicated by a yellow box that is digitally zoomed in 2B. Figure 2C shows the same area of the specimen imaged using dual-wavelength MesoTIRF, with the same ROI expanded in 2D.

**Figure 2:**
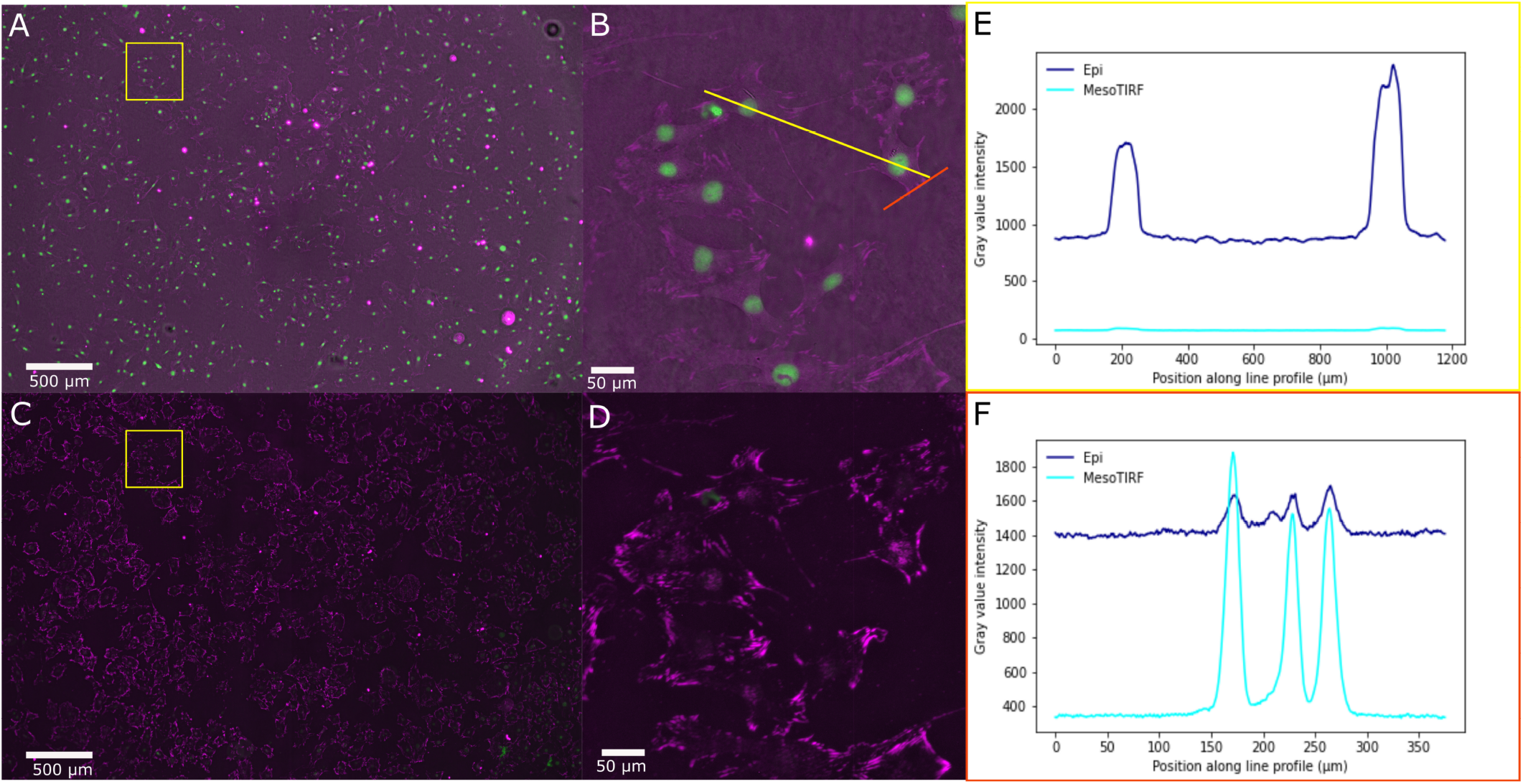
Comparison imaging of WF epi and MesoTIRF: fixed 3T3-L1 cells labelled with SYTOGreen stain visualizing nuclei (shown in green) and with an anti-paxillin antibody conjugated to Alexa Fluor Plus 594 (shown in magenta). A: WF epi image with 504 nm and 584 nm LEDs, B: ROI digital zoom of (A) with line ROIs in yellow and orange to study nuclei and paxillin respectively, C: MesoTIRF image obtained with 500 nm and 585 nm OPO SHG illumination, D: ROI digital zoom of (C), E: Yellow line profile intensity plot of neighboring nuclei in WF epi (dark blue) and MesoTIRF (cyan). F: Orange line profile intensity for three neighboring focal adhesions in WF epi (dark blue) and MesoTIRF (cyan). Fluorescently labelled nuclei are visible in WF epi data but disappear when imaged with MesoTIRF. A considerable reduction image background signal is also observed in the MesoTIRF images.

Using WF epi, the cell nuclei are clearly visible in 2B. These nuclei disappear, as expected, when imaged with MesoTIRF as shown in 2D, thus confirming that the evanescent wave in MesoTIRF is restricted to a shallow depth close to the coverslip and does not penetrate sufficiently deep into the sample to excite fluorescence from the labeled nuclei. Non-specific binding or binding of the anti-paxillin antibody to cytosolic protein is apparent when imaged with WF epi, but this fluorescence signal also disappears when using MesoTIRF illumination. This is further evidenced by the intensity profiles through two neighboring nuclei (Figure 2E) and through 3 neighboring focal adhesions (Figure 2F) for both the WF epi image (dark blue) and MesoTIRF (cyan). The difference in background is clearly evident, with MesoTIRF yielding a 4.2-fold reduction in background through the neighboring focal adhesions over WF epi, with a less noisy baseline than that of the WF epi image. While the focal adhesions are still visible in WF epi, the contrast enhancement afforded by MesoTIRF allows for the elongated features to be easily distinguished from background. An SBR improvement of 4.84 X/3.9 X/3.87 X was observed in focal adhesions 1, 2 and 3 respectively when switching from imaging with WF epi to MesoTIRF.

Furthermore, there is negligible nuclear signal in the MesoTIRF image (Figure 2E) because, as discussed previously, the 86° incident beam (resulting in a calculated evanescent field depth of 56 nm) does not penetrate the cell specimen deep enough to excite the nuclear stain. Using the ‘Surfaces’ feature in Imaris on the nuclear channel, 743 cells were counted in this single image. For the purposes of presentation, each image presented in Figure 2 has undergone the ‘Enhance Local Contrast (CLAHE)’ function in ImageJ^17^ but all analysis has been performed on raw image data.

The application of MesoTIRF for imaging of dual-labeled specimens is shown in Figure 3. Figure 3 shows a 4.4 mm x 3.0 mm FOV dual-colour MesoTIRF image with focal adhesions in magenta and F-actin in cyan. Yellow boxes show digitally zoomed images of six separate ROIs separated by a minimum distance of 0.5 mm. In all images, focal adhesions and the F-actin network adjacent to the basal cell membranes are clearly visible. We note there is a less than a 50% decrease in fluorescence signal from the centre to the edge of the imaged field, which we attribute to the Gaussian intensity profile of the illumination yielded from the optics chosen in Figure 1.

**Figure 3:**
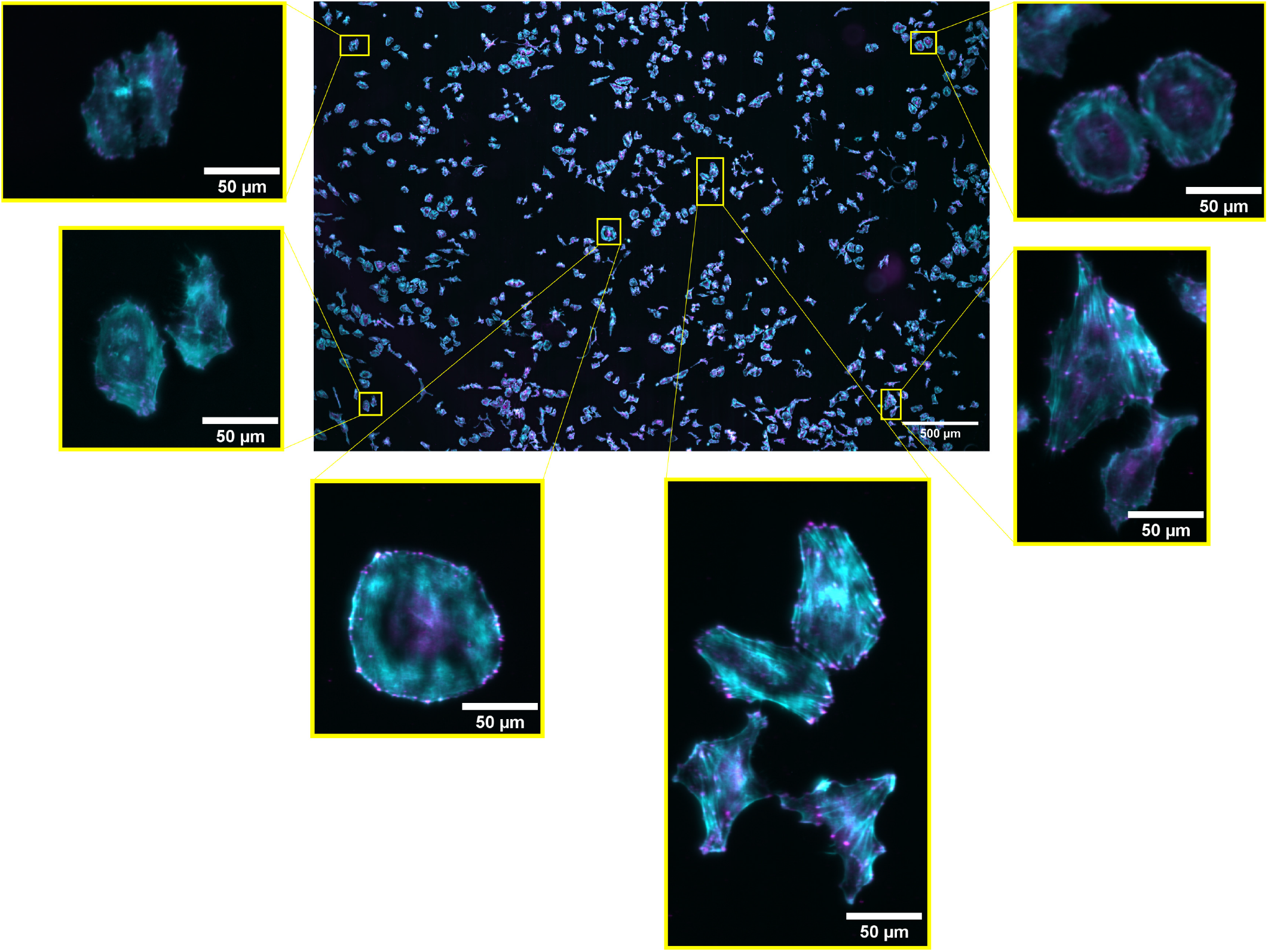
Uniformity of MesoTIRF: fixed HeLa cells labelled with an anti-paxillin antibody conjugated to Alexa Fluor Plus 594 (magenta) and Fluorescein Phallodin which stains the actin cytoskeleton (cyan). A full FOV MesoTIRF image is shown in the centre, with six ROIs indicated by yellow boxes. These show digital zoomed areas from the original dataset, and confirm a small variation in fluorescence intensity and little difference in resolvable detail across the multi-millimeter FOV.

Supplementary Figure 1 shows a further example of two-color MesoTIRF imaging with a third cell type, MeT-5A, labeled for paxillin and tubulin. We see the same improvement in image quality and contrast in this data set as demonstrated by MesoTIRF presented here, illustrating the use of the modality for a variety of cellular imaging.

## IV. DISCUSSION

We have demonstrated MesoTIRF imaging of a range of different biological samples. Coupling a custom prism TIRF illuminator with the Mesolens^12^ provides an unprecedented combination of a large FOV with sub-micron spatial resolution in three-dimensions. The optical throughput of the Mesolens is over 20 times greater than a commercial objective lens with a low magnification^12^. This presents an advantage for MesoTIRF imaging, which enables lower optical power specimen illumination with corresponding reductions in photobleaching and phototoxicity. MesoTIRF utilized the comparatively cheap prism illumination method, allowing for ease when changing evanescent field depth by varying the incidence angle of the incident beam.

In our confirmation of TIRF illumination using the specimen which was dual-labeled with a nuclear stain and an antibody against the focal adhesion protein paxillin we note that the position of the nuclei is dependent on where in their life-cycle the 3T3-L1 cells were at the point of formaldehyde fixation^20^. However we expect that the nuclear envelopes of each imaged cell is distal from the basal cell membrane, and therefore outside the reach of an evanescent field from MesoTIRF^20^.

The ability of the MesoTIRF modality to capture fine details and the contrast improvement over WF epi was examined using a fixed mammalian cell line labelled for the focal adhesion component paxillin^16^. Using the intensity signals through neighbouring focal adhesions, an average 4.2-fold improvement in SBR was measured in MesoTIRF over WF epi, with the improvement in contrast allowing for many more focal adhesions to be resolved with this novel modality (Figure 2).

The drop off in intensity from the centre to the edge of the MesoTIRF FOV is to be expected for an evanescent field generated using a Gaussian beam. This can be corrected for with flat field correction, a commonplace post-processing technique for many imaging modalities. However, as evident from the chosen ROIs in Figure 3, the level of structural detail resolvable even in these dimmer peripheral areas remains of the quality expected of TIRF.

Excitation wavelengths for MesoTIRF are presently limited to the two discussed here by the large diameter Pinkel-type custom filters used for fluorescence detection. Additional custom filters would allow this to be extended for further wavelengths.

Mesolens data are rich in information^21^ but we recognise that an imaging rate of 0.2 Hz for MesoTIRF is insufficient for several applications *in vitro*, such as cell signalling studies as reported by Crites et al^22^. However, with recent innovations in camera technologies, notably the development of cameras using large, high resolution 250 Mpixel sensors such as the Canon 2U250MRXSAA CMOS sensor, tenfold higher imaging speeds (2.4 fps) can be achieved by avoiding the need for chip shifting. In combination with environmental control, this will offer opportunities to study faster dynamic processes, for example, the action of fast-acting antimicrobial peptides^23^ or imaging of calcium transients in the plasma membrane^24^.

MesoTIRF may have applications in high-content screening^10^ or wound healing models^25^, where large cell populations must be imaged to obtain statistically significant results. However, at present MesoTIRF is only compatible with imaging at room temperature as there is no environmental imaging chamber that is compatible with the Mesolens. We are presently considering chamber designs that would be suitable for long-term imaging applications including MesoTIRF.

A present limitation of MesoTIRF is the numerical aperture of the Mesolens: at 0.47, this is much lower than a typical TIRF lens and hence the lateral resolution is around threefold poorer than a commercial objective TIRF microscope. However, with the principle of MesoTIRF now proven, an obvious next step is to add structured illumination to produce MesoTIRF-SIM^26^. Achieving SIM on the Mesolens is not a trivial task and would require either further optics in the MesoTIRF path to impose variable modulation patterns on the incident excitation beam or utilizing computational methods such as blind-SIM^27^ algorithms. This would facilitate applications in single molecule localization microscopy in cell specimens approximately two orders of magnitude larger than current technology can image. At present we are again limited by the chip-shifting camera technology, but we are carefully following developments in this field.

