## Supplementary Information for "MesoTIRF: a Total Internal Reflection Fluorescence illuminator for axial super-resolution membrane imaging at the mesoscale"

S. Foylan,[1] W. B. Amos,[2] J. Dempster,[3] L. Kölln,[2,4,5] C. G. Hansen,[4,5] M. Shaw,[6,7] and G. McConnell[1]
[1]) *University of Strathclyde, Department of Physics, Glasgow G4 0NG, UK*
[2]) *University of Strathclyde, Department of Physics, Glasgow, G4 0NG, UK*
[3]) *Strathclyde Institute for Pharmacy & Biomedical Sciences, University of Strathclyde, Glasgow, G4 0RE, UK*
[4]) *University of Edinburgh, Centre for Inflammation Research, Edinburgh, EH16 4TJ, UK*
[5]) *University of Edinburgh, Institute for Regeneration and Repair, Edinburgh, EH16 4UU, UK*
[6]) *Department of Chemical and Biological Sciences, National Physical Laboratory, Teddington, TW11 0LW, UK*
[7]) *Department of Computer Science, University College London, London, WC1E 6BT, UK*

(*Electronic mail:)

(Dated: 19 August 2022)

Here, we present the preparation protocols for cell culture, fixation and labeling of specimens prepared in the main manuscript and a further example of cell imaging with our novel MesoTIRF modality.

### I. CELL PREPARATION

The cell lines 3T3-L1*, HeLa*, and MeT-5A** were maintained in culture, at 37°C, 5% CO2 in DMEM* (11500596, Gibco) or RPMI-1640** (15-040-CV, Scientific Laboratory Supplies), 500 mL media supplemented with 50mL heat-inactivated FBS (10500064, Gibco), 5 mL Penicillin Streptomycin (P4458-100ML, Merck), 5 mL L-Glutamine (392-0441, VWR), 5 mL Sodium Pyruvate (11360-070-100ML, Gibco), 1 mL HEPES buffer (14175-095, Gibco). Coverslips were coated in a 1:500 dilution of fibronectin bovine plasma (F1171-2MG, Sigma Aldrich) in PBS (10010-023, Gibco) prior to seeding and grown in culture conditions for 24 hours (or until visible protrusions from the cell membrane to the coverslip where visible). The adherent cells were washed twice with PBS and fixed in freshly diluted 4% formaldehyde (47608-250ML-F, Sigma Aldrich). An immunofluorescence (IF) buffer solution (PBS, 2.5% FBS and 0.3% TritonX-100 (X100-100ML, Sigma Aldrich)) was applied to the fixed cells to both block non-specific binding and permeabilise the cell membranes, for 30 minutes at 37°C. This buffer was replaced by a 1:250 dilution of the desired primary antibody (paxillin: Recombinant Anti-Paxillin antibody [Y113], ab32084, AbCam) or (Paxillin Mouse anti-Human, Mouse, Rat, Clone: 5H11, MA513356, Invitrogen), (tubulin: Anti-α-Tubulin antibody, Mouse monoclonal, T6199, Merck) in the IF buffer. These coverslips where kept in a stable humidity culture dish, to prevent evaporation, for 24 hours at 4°C. The following day, the primary antibody labeled cells were washed three times with PBS before a 1:200 dilution of the appropriate secondary antibody in IF buffer was applied (Alexa Fluor 488 donkey anti-rabbit secondary, A21206 Invitrogen, Alexa Fluor 488 donkey anti-mouse secondary, ab150105, Abcam or Alexa Fluor Plus 594 goat anti-mouse secondary, A32742, Invitrogen). The coverslips where kept in light tight conditions at room temperature for 1 hour. Following three further PBS washes, the antibody labeled samples were mounted in 1% agarose with a 45 x 45 mm coverslip, to allow both for evanescent illumination through the plated coverslip and imaging through the larger one. 1% agarose was chosen over more conventional mountants such as gelvatol as the refractive index better matched that of an aqueous environment, allowing for index matching when imaging with water immersion. These samples were allowed to dry fully at 4°C in the dark before imaging. When specimens were dual labeled, the primary antibody step for both target proteins was carried out simultaneously while the fluorescent secondaries were applied sequentially, separated by PBS wash steps.

The nuclei in the 3T3-L1 cell specimens were labeled with a 1:150 dilution of Syto 16 Green ( S7578, Invitrogen) in IF buffer for 20 minutes at room temperature. This was followed by three PBS washes before mounting in 1% agarose (A4718-100G, Merck).

For fluorescent labeling of F-actin, fluorescein phalloidin (F432, ThermoFisher) was diluted 1:100 in IF buffer. Fixed, buffered and permeablized cells were incubated in this solution at room temperature in the dark for 1 hour. Cells were then removed from the staining solution, washed twice with PBS and mounted in 1% agarose.

### II. ADDITIONAL MESOTIRF CELL IMAGING

To demonstrate the wider applicability of MesoTIRF a different cell line (MeT-5A) was cultured, fixed, and labeled for tubulin and paxillin. This result (SI Figure 1) confirms the contrast enhancement and improvements in image quality synonymous with TIRF was achieved with MesoTIRF over hundreds of cells simultaneously, and is not a specimen dependent effect. The highlighted ROIs (SI Figure 1B and C) show how the punctate focal adhesions and bundles of spindled microtubules are clearly visible against a dark background in MesoTIRF. The variable density of cells across the substrate is also apparent due to the large FOV (4.4 mm x 3.0 mm) of the MesoTIRF image. Using a conventional TIRF objectives (FOV $\sim$ 50 µm x 50 µm), a comparable FOV could only be observed after extensive stitching and tiling of multiple images; a time consuming process which is obviated MesoTIRF.

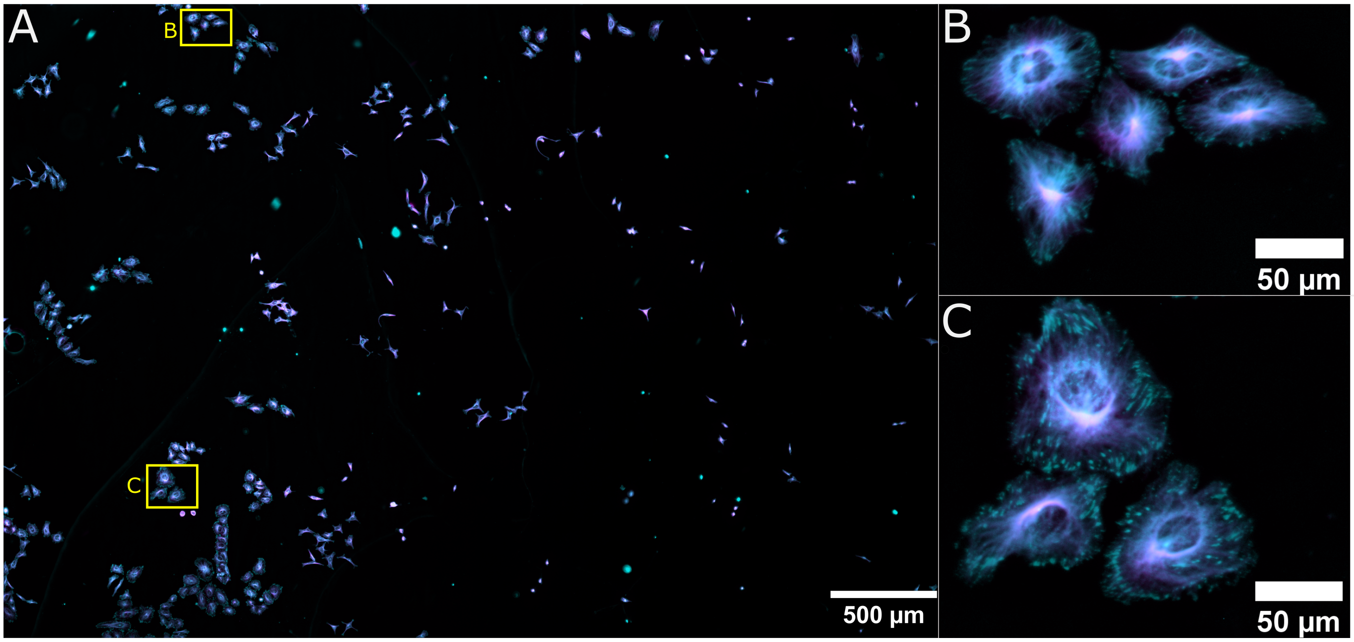


Supplementary Figure 1: MesoTIRF image of fixed MeT-5A cells labeled for tubulin with Alexa Fluor Plus 594 (magenta) and paxillin with Alexa Fluor 488 (cyan). A full FOV MesoTIRF image is shown in 1A, with two ROIs separated by several millimetres indicated by yellow boxes, and denoted by B and C. These insets B and C on the right show digital zoomed areas from the original data set, and show little change in fluorescence intensity and resolvable detail across the multi-millimetre FOV.
